# Experience-dependent plasticity modulates ongoing activity in the antennal lobe and enhances odor representations

**DOI:** 10.1101/2021.04.28.441745

**Authors:** Luis M. Franco, Emre Yaksi

## Abstract

Ongoing neural activity has been observed across several brain regions and thought to reflect the internal state of the brain. Yet, it is not fully understood how ongoing brain activity interacts with sensory experience and shape sensory representations. Here, we show that projection neurons of the fruit fly antennal lobe exhibit spatiotemporally organized ongoing activity in the absence of odor stimulation. Upon repeated exposure to odors, we observe a gradual and long-lasting decrease in the amplitude and frequency of spontaneous calcium events, as well as a reorganization of correlations between olfactory glomeruli during ongoing activity. Accompanying these plastic changes, we find that repeated odor experience reduces trial-to-trial variability and enhances the specificity of odor representations. Our results reveal a previously undescribed experience-dependent plasticity of ongoing and sensory driven activity at peripheral levels of the fruit fly olfactory system.

## INTRODUCTION

Neural circuits display continuous fluctuations in their activity, even in the absence of sensory stimuli or environmental alterations. This ongoing neural activity is widespread in the brain, spanning from sensory systems to higher brain regions, and it has been associated with various cognitive functions (Raichle, 2010; Sadaghiani et al., 2010). Despite being previously considered noise, ongoing neural activity is highly structured, often exhibiting spatiotemporally organized patterns (Arieli et al., 1996; Bartoszek et al., 2021; Fox et al., 2005; Jetti et al., 2014; Marques et al., 2020; Romano et al., 2015; Shimaoka et al., 2019), reflecting the functional connectivity and internal states of neural networks (Andalman et al., 2019; Lange and Haefner, 2017; Lovett-Barron et al., 2017). Moreover, accumulating evidence indicates that continuous interactions between externally-driven sensory representations and internally-generated ongoing activity shape each other. For instance, during specific tasks, ongoing neural activity is transiently suppressed in a large area of the human cortex, except in regions engaged in the task (Raichle, 2010; Raichle et al., 2001). Similarly, sensory-driven transient transformations of ongoing activity have been observed in several brain regions across different species (Galán et al., 2006a; Hahn et al., 2012; Ichinose et al., 2017; Romano et al., 2015; Shimaoka et al., 2019; Vanni and Murphy, 2014; Wosniack et al., 2021). Interestingly, repeated exposure to sensory stimuli can also lead to long term experience-dependent alterations in neural circuit connectivity and activity. Such experience-dependent alterations have been shown to stabilize neuronal responses to behaviorally relevant sensory stimuli and even enhance stimulus discriminability (Bazhenov et al., 2005; Bhandawat et al., 2007; Galán et al., 2006b; Jacobson et al., 2018; Kilgard and Merzenich, 1998; Musall et al., 2019; Poort et al., 2015; Stopfer and Laurent, 1999).

In the olfactory system, ongoing activity in the absence of odors is reported from the level of olfactory sensory neurons (Bhandawat et al., 2007; Friedrich and Laurent, 2004; Nagel et al., 2015; Olsen et al., 2010) to the olfactory bulbs (Fujimoto et al., 2019; Galán et al., 2006b; Gorin et al., 2016; Nagel and Wilson, 2016; Padmanabhan and Urban, 2010) and the olfactory cortex (Lottem et al., 2016; Popov and Szyszka, 2020; Rojas-Líbano et al., 2014; Wilson and Yan, 2010). To some extent, this ongoing activity can be interpreted as background noise, which is generated by the internal biophysical properties of neurons at each level (Kazama and Wilson, 2009; Padmanabhan and Urban, 2010). However, evidence from the vertebrate olfactory bulb (Gorin et al., 2016; Padmanabhan and Urban, 2010) and the insect antennal lobe (Galán et al., 2006b) suggests that ongoing neural activity reflects the structured synchronous activity of neuronal ensembles in the olfactory system. Moreover, top-down modulation of olfactory bulb inhibitory interneurons is also known to modulate not only sensory responses but also the resting state of ongoing activity in olfactory circuits (Bundschuh et al., 2012; Dacks et al., 2009; Liu, 2020; Lottem et al., 2016; Petzold et al., 2009). In fact, sensory experience is known to play an important role in modulating olfactory representations and synchrony across multiple levels of the olfactory pathway (Galán et al., 2006b; Jacobson et al., 2018; Rojas-Líbano et al., 2014). Yet, the link between experience-dependent plasticity of odor representations and ongoing neural activity is not fully understood.

In this study, we showed that the projection neurons in the fly antennal lobe exhibit spatiotemporally organized ongoing activity in the absence of odor stimulation. Repeated exposure to odors reduces the amplitude and frequency of spontaneous calcium events and alters correlations between olfactory glomeruli during ongoing activity. Moreover, this experience-dependent plasticity decrease trial to trial variability of odor representations, thereby increasing the robustness and specificity of olfactory computations.

## RESULTS

To investigate the spatial and temporal features of ongoing activity in Drosophila antennal lobe, we measured spontaneously generated calcium signals in GH146-Gal4;UAS-GCaMP6m fruit flies expressing the calcium indicator GCaMP6m in the projection neurons of the antennal lobe (Fig. 1A). To evaluate the impact of repeated odor exposure on the ongoing activity of the antennal lobe, we applied 3 trains of 8 consecutive stimuli with the same odor, interleaved by epochs of 6 minutes long ongoing activity (Fig. 1A-C). This experimental design was tested using 8 different odors (1-pentanol, 2-heptanone, 3-octanol, 4-methylcyclohexanol, amyl acetate, benzaldehyde, ethyl acetate and ethyl valerate) in different experiments. Individual olfactory glomeruli were identified (Fig. 1B,C) by using a previously described independent component analysis (Franco et al., 2017; Mukamel et al., 2009). Altogether, we observed that the number and the amplitude of spontaneously occurring calcium events decreased significantly during every epoch after exposure to repeated odor stimulation (Fig. 1D-F). Intriguingly, this overall reduction of ongoing glomerular activity was also accompanied by a redistribution of pairwise correlations between individual glomeruli, indicating a functional reorganization of the antennal lobe. We also observed that functional clusters of olfactory glomeruli were stable across all epochs of ongoing activity, where pairs of olfactory glomeruli remained in their respective k-means clusters (Fig. S1). These results indicate a significant, gradual and lasting experience-dependent modulation of the ongoing activity in Drosophila antennal lobe circuits upon repeated odor exposure.

**Figure 1.**
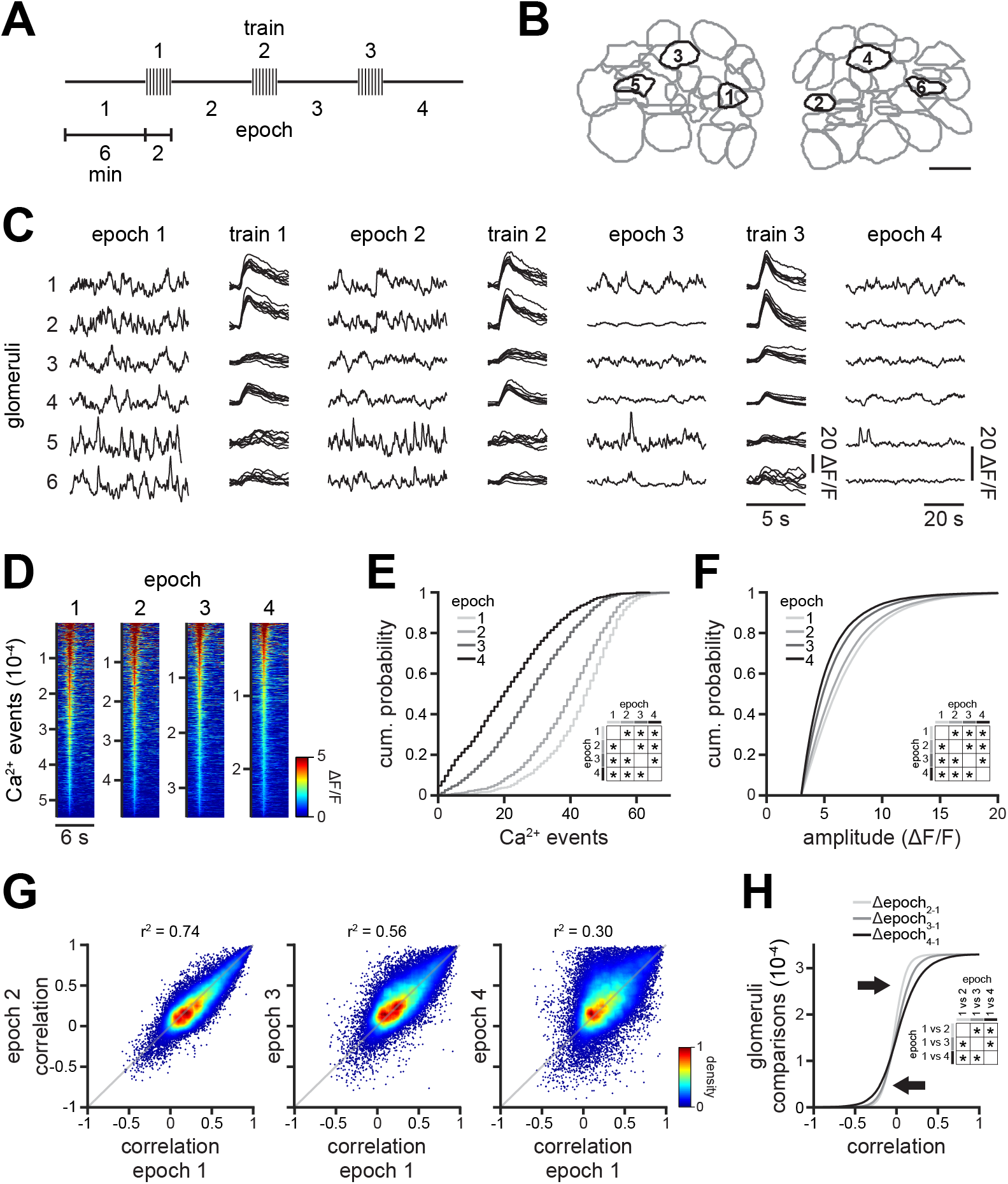
Repetitive odor exposure dampens ongoing activity in the antennal lobe. A. Experimental design consisting of three trains of eight odor stimuli interleaved by four epochs of ongoing activity. 8 different odors were tested in parallel experiments, see methods for details. B-C. Representative example of glomerular locations (B) and their corresponding ongoing activity epochs and average responses during odor stimulation trains (C). Scale bar, 10 μm. D. Heat maps showing all individual calcium events across four ongoing activity epochs. E-F. The number of calcium events (E; n = 1,268 glomeruli in 24 flies; Wilcoxon rank-sum test; epoch 1 vs epoch 2, p = 3.7 x 10-20; epoch 1 vs epoch 3, p = 2.7 x 10-159; epoch 1 vs epoch 4, p = 5.2 x 10-246) as well as their amplitude (F; n = 1,268 glomeruli in 24 flies; Wilcoxon rank-sum test; epoch 1 vs epoch 2, p = 1.2 x 10-97; epoch 1 vs epoch 3, p = 0; epoch 1 vs epoch 4, p = 0) decreased with repeated odor stimulation. G. Pairwise Pearson’s correlations of ongoing activity among all glomeruli compared across epochs. Correlations of ongoing glomerular activity changes upon repeated odor stimulation (n = 32,947 glomerulus pairs). H. Cumulative distribution of the change in correlation coefficients for all recorded glomeruli, during 4 different epochs. The distribution of the change in pairwise correlations among glomeruli significantly changed by repetitive odor stimulation (n = 32,947 glomerulus pairs; Kolmogorov-Smirnov test; Δepoch2–1 vs Δepoch3–1, p = 7.8 x 10-235; Δepoch2–1 vs Δepoch4–1, p = 0, Δepoch3–1 vs Δepoch4–1, p = 1.3 x 10-155). Arrows indicate the redistribution of glomeruli correlations, wherein some glomeruli become more positively or negatively correlated.

Next, we asked whether each train of odor exposure, modulating ongoing glomerular activity, can also lead to gradual and lasting alterations in odor representations. We observed a gradual and cumulative reduction of odor responses upon exposure to each train of odor stimuli (Fig. 2A-C, S2A-B). To quantify whether this experience-dependent reduction in odor responses is different across olfactory glomeruli, we categorize olfactory glomeruli based on their response amplitude. We find that repetitive odor exposure significantly increases the number of non-responsive (or silent) glomeruli (Fig. 3D-E, shown in blue) and reduces the number of odor responsive (or active) glomeruli (Fig. 3D-E, shown in green, yellow and red).We hypothesized that such reorganization of glomerular response amplitudes may lead to an experience-dependent sparsening of odor representation, where repeated exposure to odors gradually silence most glomeruli, except few specific and strongly responding glomeruli. To quantify this, we used the measure of population sparseness, which would be equal to 1 only if a single specific glomerulus responds to an odor, and would be equal to 0 if all glomeruli equally respond to a given odor. In fact, we observed a significant, gradual and long-lasting increase in population sparseness upon each exposure to the trains of odor stimuli (Fig. 2F).

**Figure 2.**
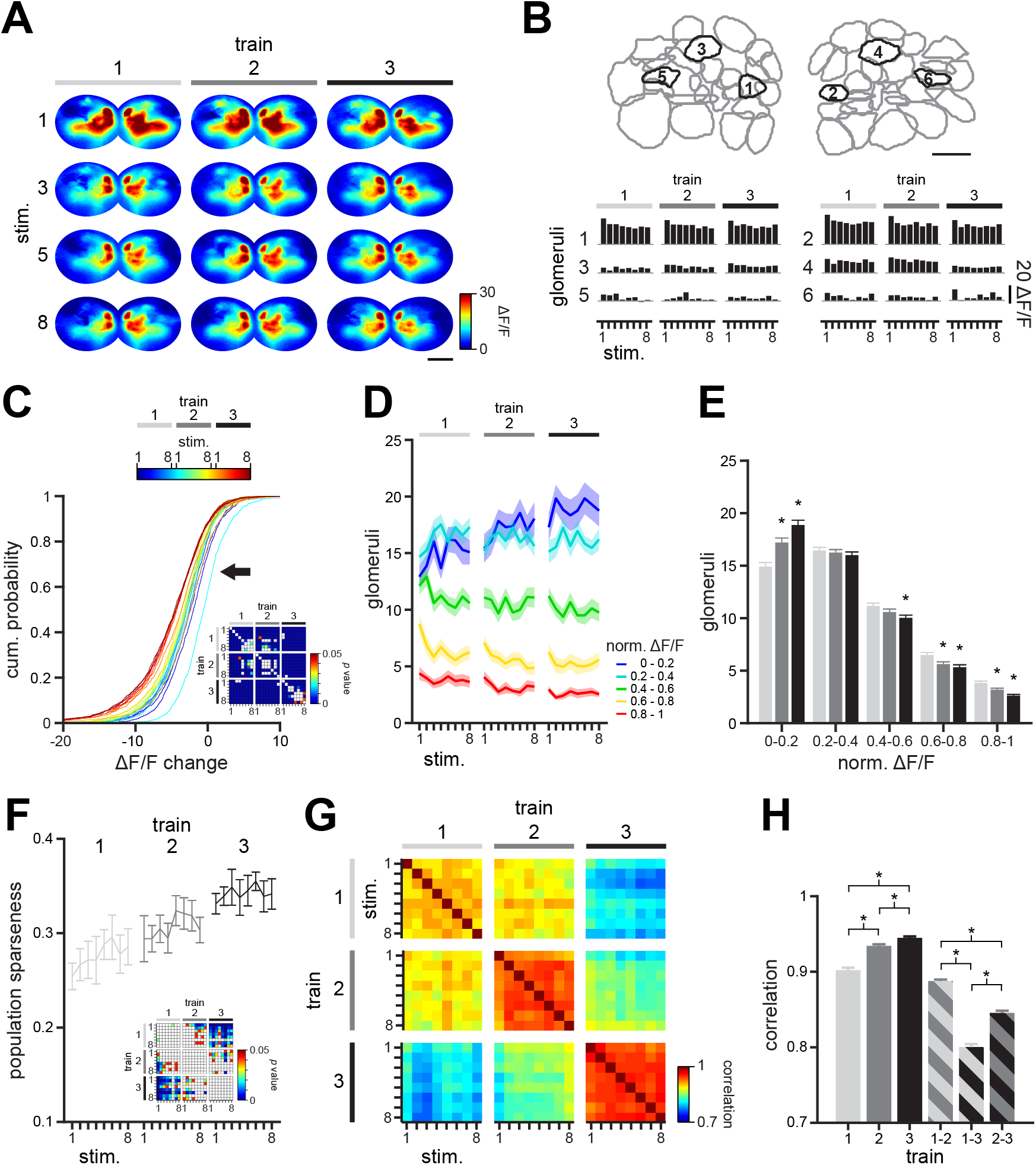
Repetitive odor stimulation increases reliability of odor representations. A. Representative odor maps calculated by the percent change of fluorescence intensity (ΔF/F) during 1.5 s after response onset. Warmer colors signify strong responses. Scale bar, 20 μm. B. Examples of the location of glomeruli (top) and their corresponding odor-evoked excitation and inhibition responses (bottom). Scale bar, 10 μm. C. Cumulative distribution of the change in odor responses, compare to the first odor response. Repetitive stimulation progressively and significantly decreases glomerular responses (n = 1,268 glomeruli in 24 flies across 3 trains of 8 stimuli each, Wilcoxon signed-rank test, p values lower than 0.05 are color coded for each comparison in the inset). The arrow indicates a displacement of the cumulative distribution of odor responses to the left, indicating a reduction in odor responses. D. Number of glomeruli grouped by their relative response magnitude across the three trains of stimuli. Note that whereas the number of odor responsive glomeruli (red) decreases, the number of weak or non-responsive glomeruli (blue) significantly increases (see panel E) with repetitive stimulation. (mean ± s.e.m., n = 1,268 glomeruli in 24 flies). E. Bar plots showing the average number of glomeruli grouped by their magnitude for each stimulus train. The number of weak or non-responsive glomeruli significantly increases with repetitive stimulation, whereas the number of responsive glomeruli significantly decrease (mean ± s.e.m., n = 1,268 glomeruli in 24 flies; Wilcoxon rank-sum test; 0-0.2 ΔF/F, train 1 vs train 2, p = 5.4 x 10-5; train 1 vs train 3, p = 7.7 x 10-11; 0.2-0.4 ΔF/F, train 1 vs train 2, p = 0.74; train 1 vs train 3, p = 0.30; 0.4-0.6 ΔF/F, train 1 vs train 2, p < 0.14; train 1 vs train 3, p = 5.9 x 10-3; 0.6-0.8 ΔF/F, train 1 vs train 2, p < 0.02; train 1 vs train 3, p = 6.3 x 10-4; 0.8-1.0 ΔF/F, train 1 vs train 2, p < 2.0 x 10-3; train 1 vs train 3, p = 1.6 x 10-8). F. Population sparseness of odor representations across stimulus trains. Sparseness increases with repetitive stimulation, indicating more specific recruitment of a small number of glomeruli upon odor stimulation (mean ± s.e.m., n = 24 flies exposed to 8 different odors in 3 independent experimental sessions each, Wilcoxon rank-sum test, p values lower than 0.05 are color coded for each comparison in the inset). G. Matrix representing the average pairwise correlations among odor representations elicited in the three trains of odor stimuli (n = 24 flies exposed to 8 different odors in 3 independent experimental sessions each; odor representations were calculated for each experimental session and compared across the 3 trains of 8 stimuli). H. Correlations among odor representations within and across stimulus trains. Representations become significantly more reliable after repetitive stimulation (mean ± s.e.m., n = 24 flies exposed to 8 different odors in 3 independent experimental sessions each; Wilcoxon rank-sum test; within train 1 vs within train 2, p = 8.7 x 10-15; within train 1 vs within train 3, p = 3.7 x 10-28; within train 2 vs within train 3, p = 2.9 x 10-5; across trains 1 and 2 vs across trains 1 and 3, p = 2.7 x 10-68; across trains 1 and 3 vs across trains 2 and 3, p = 1.6 x 10-16; across trains 1 and 2 vs across trains 2 and 3, p = 1.2 x 10-24).

**Figure 3.**
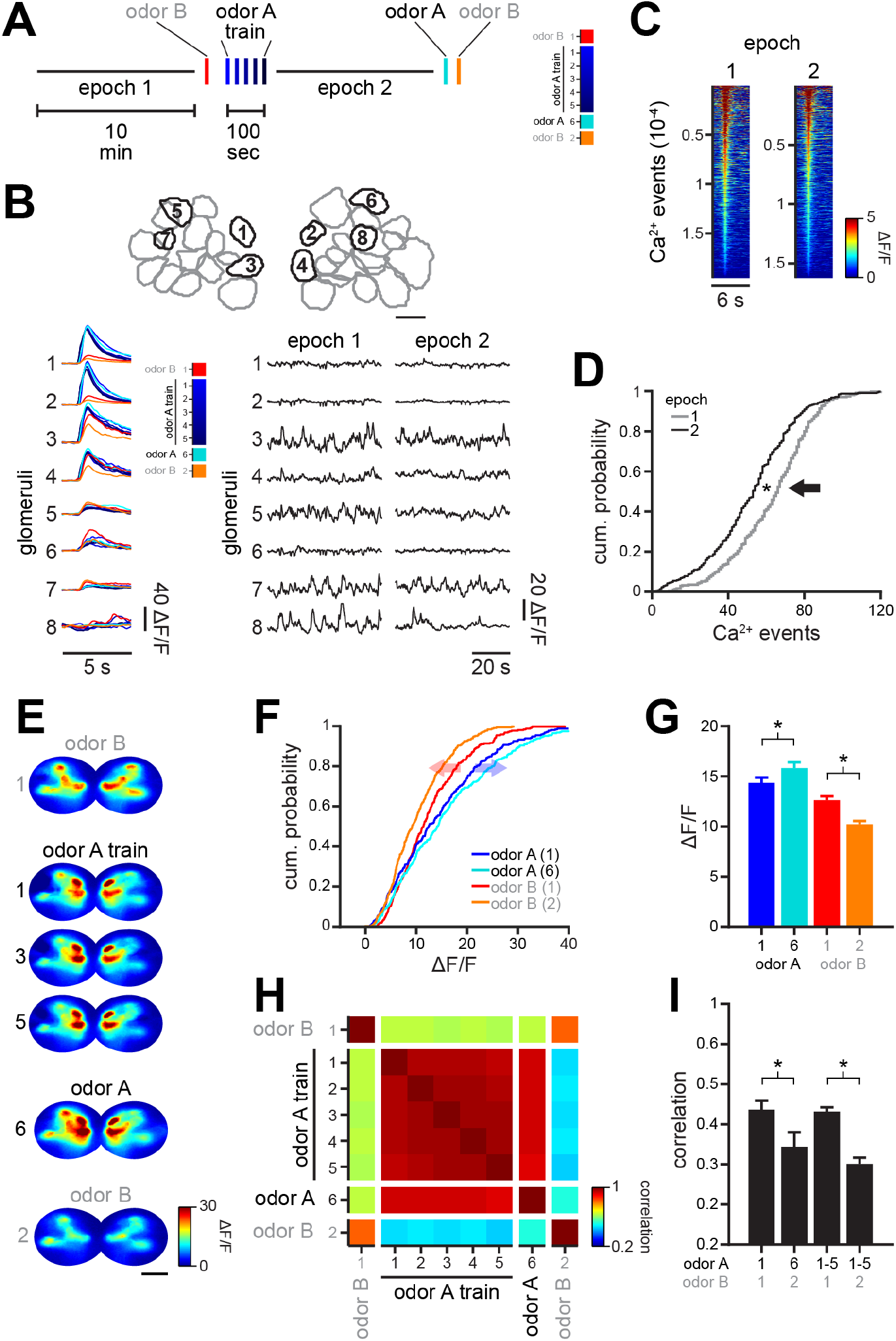
Repetitive odor exposure facilities discrimination of odor representations. A. Experimental design consisting of one train of five odor stimuli (blue) preceded by one stimulus with a naïve odor (red) interleaved by epochs of ongoing activity. After the entrainment of the antennal lobe and 10 min waiting period, both odors are presented again at the end of the experiment (cyan and orange). B. Examples of the location of glomeruli (B) and their corresponding odor-evoked responses (colored traces), and ongoing activity epochs. Scale bar, 10 μm. C. Heat maps showing all of the detected calcium events across the two ongoing activity epochs. D. Cumulative distribution of the number of calcium events. Note that the number of calcium events decrease with repetitive stimulation (n = 309 glomeruli in 10 flies, Wilcoxon rank-sum test, p = 3.3 x 10-9). E. Odor maps calculated by the percent change of fluorescence intensity (ΔF/F) during 1 s after response onset. Warmer colors signify strong responses. Scale bar, 20 μm. F. Cumulative distribution of all glomerular responses for the first and last exposure to odor A and B, respectively (n = 309 glomeruli in 10 flies). Note that responses to the unrepeated odor B decrease (red arrow), whereas responses to the repeated odor A increase (blue arrow). G. Odor responses to the repeated odor A significantly increased after repetitive stimulation, whereas responses to the unrepeated odor B significantly decreased (n = 309 glomeruli in 10 flies; Wilcoxon signed-sum test; odor A1 vs odor A6, p = 4.9 x 10-11; odor B1 vs odor B2, p = 1.3 x 10-19).

We expect that an experience-dependent sparsening of odor responses could in principle lead to more robust odor representations with reduced trail-to-trial variability. To test this hypothesis, we calculated pairwise similarities of repeated odor representations by calculating Pearson’s correlations (Fig. 2G) and Euclidean distances (Fig. S2C) of glomerular response patterns. Both of these comparisons revealed a significant decrease in trial-to-trial variability (Fig. 2H, S2D), enhancing the robustness of odor representations up on repeated odor exposure. Altogether, our results indicate that the antennal lobe circuit gradually undergoes experience-dependent plasticity upon persistent odor exposure, reducing neural noise and trial-to-trial variability, and thereby improving the specificity of odor representations.

One possible consequence of the observed experienced-dependent increase in the specificity of odor representations, is a potential improvement of odor discriminability. To test this hypothesis, we compared the odor response patterns of two different odors, before and after exposing the antennal lobe circuits to repeated stimulation with only one of the odors (Fig. 3A). Similar to our results with multiple odor train stimulations, we observed that even a single odor train with 5 repetitions lead to alterations in both glomerular odor responses and ongoing activity (Fig. 3B). After 5 odor repetitions we observed a decrease in the number of spontaneous calcium events (Fig. 3C-D) as well as in correlations in ongoing glomerular activity (Fig. S3A-B). Accompanying these experience-dependent changes, we observed that glomerular responses to the repeated odor became more different from the responses to the unrepeated odor, even after 10 min of ongoing activity period with no odor stimulation (Fig. 3E-G). We compared the similarities of odor representations of these two odors, before and after repeated exposure to one of the odors, by using Pearson’s correlations (Fig. 3H-I) and Euclidean distances (Fig. S3C-D). All analysis confirmed that the representation of two different odors become more dissimilar, and thus easier to discriminate upon repeated exposure to only one of the odors. Together, these results show that experience-dependent alterations in the antennal lobe improve odor discriminability upon repeated odor exposure.

## DISCUSSION

In this study, we showed that the projection neurons of the fly antennal lobe display structured ongoing activity. We also showed that repeated odor-exposure triggers experience-dependent plasticity in the antennal lobe, decreasing ongoing activity, which in turn facilitates more robust odor representations with reduced trial to trial variability and increased odor specificity. These findings are particularly relevantfor understanding neural computations underlying olfactory-guided behaviors. When animals navigate across odor plumes, they often encounter not only a single continuous odor pulse, but rather experience repeated odor pulses with dynamically changing exposure frequency (Boie et al., 2018; Van Breugel and Dickinson, 2014; Lewis et al., 2020; Louis et al., 2008; Park et al., 2016; Porter et al., 2007; Reiten et al., 2017; Vickers et al., 2001; Victor et al., 2019). Hence, such repeated odor exposure, as we investigated in this study, is likely a common way for animals to experience odor plumes in nature. We argue that the experience-dependent enhancement of odor representations upon repeated exposure to the same odor, may serve an important function during odor guided navigation by adjusting internally generated dynamics of the antennal lobe circuit.

We observed that odor experience-dependent plasticity dampens ongoing activity fluctuations and reduce the internal noise of the antennal lobe. These alterations are accompanied by an increase in the number of silent glomeruli and broadly reduce the amplitude of glomerular odor responses. A potential mechanism for such plasticity may rely on a gradual recruitment of the inhibitory local interneurons of the antennal lobe. A relatively large fraction of antennal lobe local interneurons (LNs) are inhibitory (Berck et al., 2016; Chou et al., 2010; Nagel et al., 2015; Olsen et al., 2010; Sachse et al., 2007), regulating the activity of both projection neurons (Franco et al., 2017; Grabe et al., 2020; Wilson and Laurent, 2005) and olfactory receptor neuron terminals (Olsen and Wilson, 2008; Root et al., 2008). Thus, it is likely that repeated odor exposure gradually increases the activity of inhibitory LNs, leading to accumulating levels of the inhibitory neurotransmitter GABA release in the antennal lobe. Such recruitment of inhibitory neurons have been previously shown during odor habituation (Das et al., 2011). Our results suggested that this excessive inhibition might in turn lead to a persistent dampening of ongoing antennal lobe activity, reducing noise levels for extended periods of time, beyond the transient odor exposure. Importantly, such inhibition, mediated by local interneurons, is preserved across the vertebrate olfactory bulb (Abraham et al., 2010; Arevian et al., 2008; Tabor et al., 2008).

Recent studies have demonstrated strong and lasting neuromodulatory control of inhibitory interneurons in the vertebrate olfactory bulb via dopaminergic (Banerjee et al., 2015; Bundschuh et al., 2012; Hsia et al., 1999) and serotonergic (Brill et al., 2016; Dugué and Mainen, 2009; Petzold et al., 2009) modulation. Interestingly, the Drosophila neuromodulator octopamine is known to regulate inhibitory interneurons in the antennal lobe (Rein et al., 2013) and to influence the internal state of the fly during odor-guided behaviors, such as searching for appropriate food sources (Christiaens et al., 2014) depending on the current nutritional demands (Corrales-Carvajal et al., 2016). Hence, it is also possible that repeated odor exposure entrains these neuromodulatory systems, which can lead to lasting and more effective inhibition in the olfactory bulb and antennal lobe circuits, reducing noise levels and enhancing the specificity of odor representations. In consonance with this, it was recently reported that pre-exposure to odors recruits dopaminergic neurons, producing latent inhibition in the mushroom body, which leads to changes in appetitive behavior in the fruit fly (Jacob et al., 2021). Thus, neuromodulatory systems help shaping odor representations through local inhibition at different stages in the olfactory processing pathway. However, further research is needed in order to elucidate the mechanisms by which inhibitory interneurons and neuromodulatory systems are recruited upon repeated odor exposure.

We argue that the sensory experience-dependent plasticity that we observe in the antennal lobe represents a phenomenon that can be translated to the rest of the brain. This is mainly because such sensory induced dampening of the ongoing neural activity is prominent in many parts of the central nervous system (Raichle, 2010; Raichle et al., 2001). Hence, sensory experience-dependent dampening of the internal noise of neural networks can contribute to more effective and more specific processing of incoming information. Investigating the presence and mechanisms of this phenomenon is important for understanding how higher brain regions, such as the habenula (Baker et al., 2015; Jetti et al., 2014; Mizumori and Baker, 2017), thalamus (Kirouac, 2015; MacLean et al., 2005) or cortex (Blaeser et al., 2017; Tan, 2015), contribute to the switching of brain states. Our results revealed that experience-dependent alterations are not only restricted to higher brain regions, but can specifically modulate sensory representations at the first stage of sensory pathways, such as the antennal lobe.

## ACKNOWLEDGEMENTS

We thank Ilona Kadow (who shared UAS-GCaMP6m flies) and Rachel Wilson (who shared GH146-GAL4 flies). This work was funded by VIB (E.Y.), NERF (E.Y.), the VIB International PhD Program (L.M.F.) and ERC Starting Grant 335561 (E.Y). The E.Y. lab is currently funded by the Kavli Institute for Systems Neuroscience at NTNU.

## AUTHOR CONTRIBUTIONS

Conceptualization, L.M.F. and E.Y.

Methodology, data, analysis L.M.F.

Writing, L.M.F. and E.Y.

Funding acquisition, L.M.F. and E.Y.

Supervision, E.Y.

The authors declare no competing interests.

## MATERIALS AND METHODS

### Olfactory stimulation

Odors for protocol 1 (Fig. 1,2,S1,S2) are: 1-pentanol, 2-heptanone, 3-octanol, 4-methylcyclohexanol, amyl acetate, benzaldehyde, ethyl acetate and ethyl valerate. Odors for protocol 2 (Fig. 3,S3) are: ethyl acetate (repeated) and benzaldehyde (unrepeated). All odors were diluted 1:100 v/v in paraffin oil. Odors were delivered through a custom-built mechanism, which dilutes the headspace of the odor vial a further 10-fold in clean air. The flow rate of odor delivery was 2.2 L/min. Odors were applied for 500 ms, either 8 times (protocol 1) or 5 times (protocol 2) per train of stimuli, with an inter-trial interval of 15 s. Each train of stimuli was spaced by 6 min (protocol 1) or 10 min (protocol 2) where no odor stimulation was applied. The stimulation protocol was controlled by a Master-8 stimulator (A.M.P.I.), programmed for each protocol, respectively.

### Fly preparation and calcium imaging

Calcium imaging experiments were conducted on 5-10 days post-eclosion female GH146-GAL4/UAS-GCaMP6m;+/+ flies. For each experiment, a fly was secured to a custom chamber made of aluminum foil, smoothly shaped to fit the thorax and the upper part of the head. The fly was then fixed to the chamber with wax, surrounding the space between the fly and the top of the chamber. Also, the front and middle legs as well as the proboscis were carefully immobilized with small drops of wax to the bottom of the chamber. Next, the cuticle above the antennal lobes was carefully removed using a very sharp needle made of tungsten wire. The dorsal side of the antennal lobes was then imaged while constantly perfusing the brain with oxygenated saline. The saline contained (in mM): 103 NaCl, 3 KCl, 1 NaH2PO4, 1.5 CaCl2, 4 MgCl2, 26 NaHCO3, 5 TES, 10 glucose and 8 trehalose. GCaMP6m fluoresce was imaged at 10 Hz using an EMCCD camera (Hamamatsu Photonics) installed on an Olympus BX51 fluorescence microscope (Olympus Corporation).

### Data analysis

Fluorescence imaging stacks were processed using custom code written in Matlab (MathWorks). Briefly, independent component analysis [41,42] was performed on raw fluorescence stacks corresponding to the left and to right antennal lobes, respectively, for each experimental session. The resulting independent components were then manually curated to eliminate noisy components corresponding to either artifacts in the surrounding of the antennal lobes, or very small regions, likely arising from noisy pixels. This procedure yields a map containing clusters of pixels with similar fluorescence changes, corresponding to functional glomeruli (Fig. 1B).

To obtain the activity time series for each glomerulus, we first processed raw fluorescence stacks on a pixel-by-pixel basis, defining F0 as the average fluorescence over a 30 s sliding window, which was then used to calculate ΔF/F for each time bin. Next, we averaged the activity time series for all pixels contained in the region described by each corresponding glomerulus. The resulting glomerulus time series were used for all subsequent analyses.

Ongoing activity across glomeruli was compared by computing the Pearson’s correlation coefficient on the entire 6 min (protocol 1) or 10 min (protocol 2) traces for all glomeruli pairs and all epoch pairs corresponding to each experimental session. Similarly, k-mean clustering was performed on the full ongoing activity traces from each epoch. When assigning glomeruli into clusters, the distances between individual glomerulus were calculated based on the cosine distances between their ongoing activity. Cluster fidelity was measured as the probability of a particular glomerulus staying in the same cluster for different epochs. As control, we shuffled cluster identity of individual glomeruli and performed the calculation again.

Odor responses were defined as the average activity during a 1.5 s window after response onset for each glomerulus. To evaluate the similarity across activity patterns elicited in the antennal lobe by odor stimulation, we constructed odor vectors consisting of odor responses from all glomeruli for each experimental session. We then used the Pearson’s correlation coefficient and Euclidean distances to compare across odor vectors. In addition, we calculated the population sparseness as:

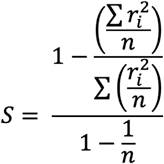

where r_i is the response of the i ^th glomerulus to a particular odor and n is the total number of glomeruli per fly. Values near 0 indicate low selectivity whereas values near 1 indicate high selectivity of the antennal lobe circuit for a particular odor. Normalized odor responses were obtained by subtracting the minimum response and then dividing by the maximum response within each fly per stimulus. We then grouped responses according to their magnitude in 5 groups: 0-0.2 ΔF/F, 0.2-0.4 ΔF/F, 0.4-0.6 ΔF/F, 0.6-0.8 ΔF/F and 0.8-1.0 ΔF/F.

All data groups were compared using non-parametric tests (Wilcoxon ranksum/signed-rank tests or Kolmogorov-Smirnov test).

## SUPPLEMENTAL FIGURES

**Figure S1. Related to Figure 1.**
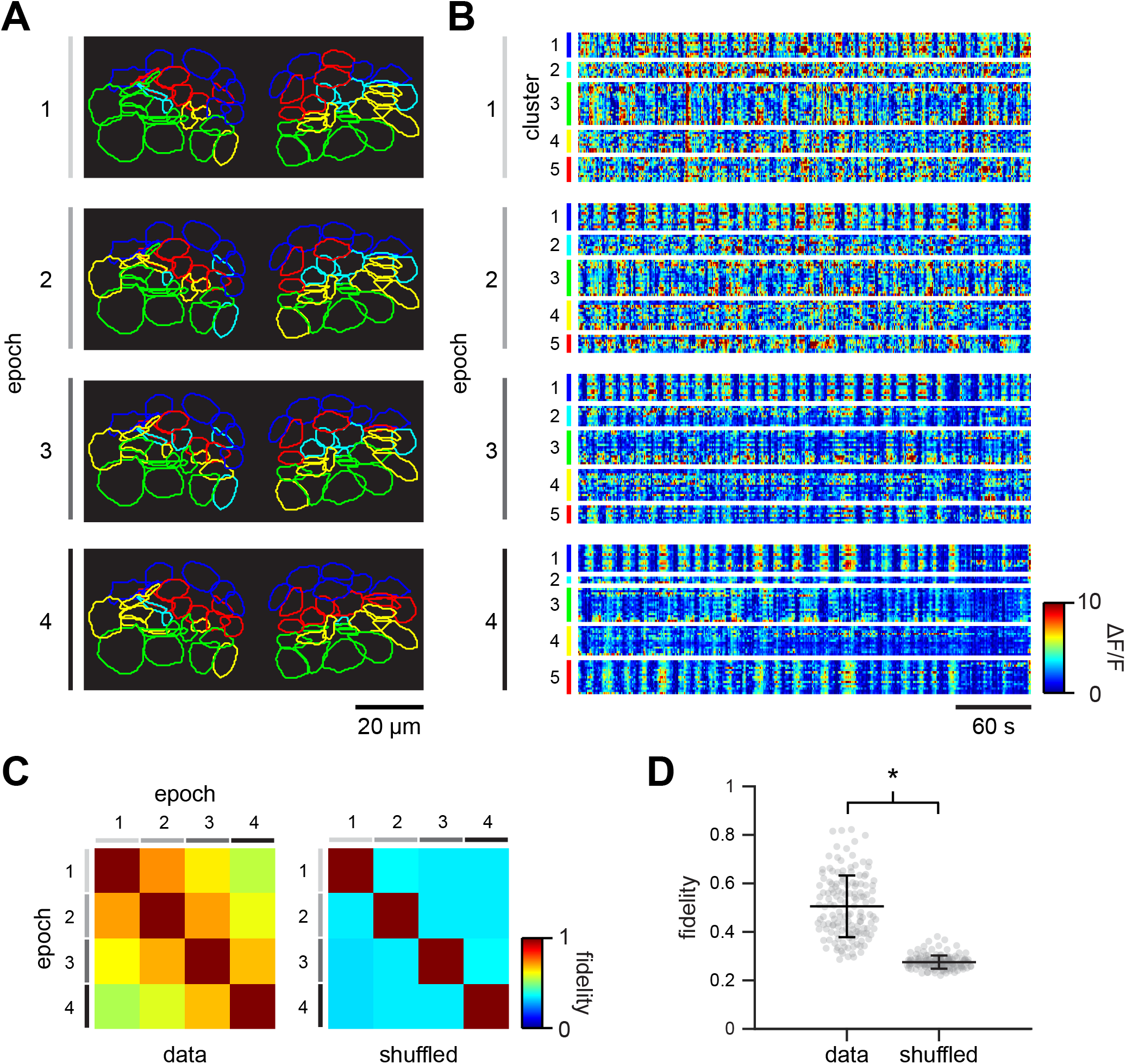
A. Functional clusters obtained by k-means of an example fly for the four ongoing activity epochs. B. Representative examples pf ongoing activity of the functional clusters shown in the example fly in A. C. Matrices showing the average cluster fidelity of neurons compared epoch by epoch (n = 24 flies). D. Average fidelity of glomerular clusters and expected chance levels for all of the functional clusters in all of the flies (mean ± s.d., n = 6 cross-epoch comparisons in 24 flies, Wilcoxon signed-rank test, p = 2.2 x 10-25). H. Matrix representing the average pairwise correlations among odor representations elicited by the repeated odor A, and unrepeated odor B (n = 10 flies). I. Correlations among odor representations for the repeated odor A and the unrepeated odor B. Note that the representations of the repeated odor A become more different from that of the unrepeated odor B (mean ± s.e.m., n = 10 flies; Wilcoxon signed-sum test; odor A1-odor B1 vs odor A6-odor B2, p = 0.02; odor A1-5-odor B1 vs odor A1-5-odor B2, p = 0.03).

**Figure S2. Related to Figure 2.**
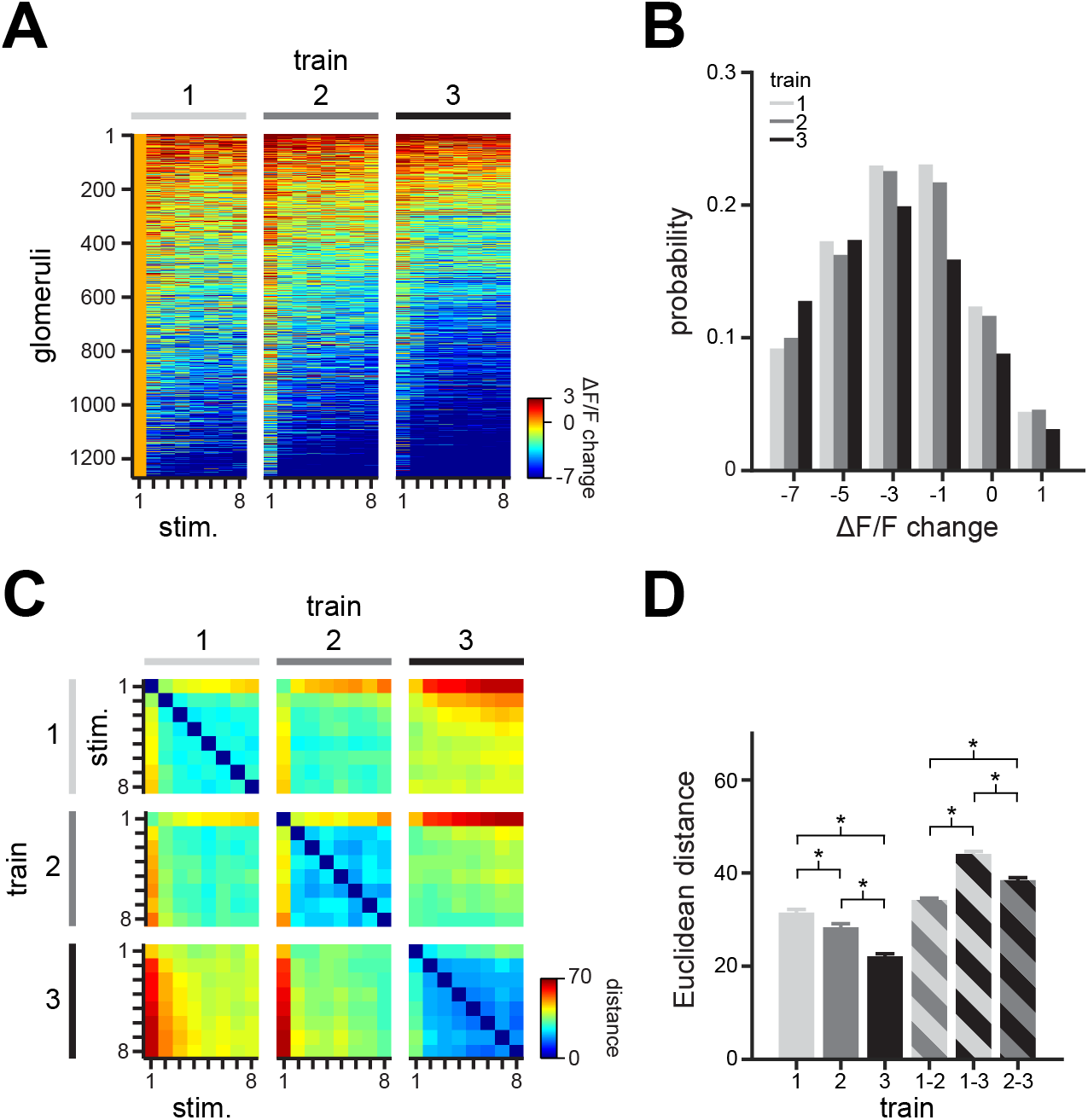
A. Odor response change from the first odor stimulus of all individual glomeruli (n = 1,268 glomeruli in 24 flies). Glomeruli are sorted based on their odor activity level, from the more responsive (top) to the less responsive (bottom). B. Histogram showing the change in glomerular responses grouped by train of stimulation (n = 1,268 glomeruli in 24 flies). Note that repetitive stimulation increases the probability of glomeruli with reduced responses and decreases the probability of glomeruli with increased responses. C. Matrix representing the average Euclidean distances among odor representations elicited in the three trains of stimuli (n = 24 flies exposed to 8 different odors in 3 independent experimental sessions each; odor representations were calculated for each experimental session and compared across the 3 trains of 8 stimuli). D. Euclidean distances among odor representations within and across stimulus trains. Representations become significantly more reliable after repetitive stimulation (mean ± s.e.m., n = 24 flies exposed to 8 different odors in 3 independent experimental sessions each; odor representations were calculated for each experimental session and compared across the 3 trains of 8 stimuli; Wilcoxon rank-sum test; within train 1 vs within train 2, p = 1.3 x 10-6; within train 1 vs within train 3, p = 1.4 x 10-26; within train 2 vs within train 3, p = 7.5 x 10-10; across trains 1 and 2 vs across trains 1 and 3, p = 1.0 x 10-45; across trains 1 and 3 vs across trains 2 and 3, p = 3.6 x 10-19; across trains 1 and 2 vs across trains 2 and 3, p = 1.4 x 10-9).

**Figure S3. Related to Figure 3.**
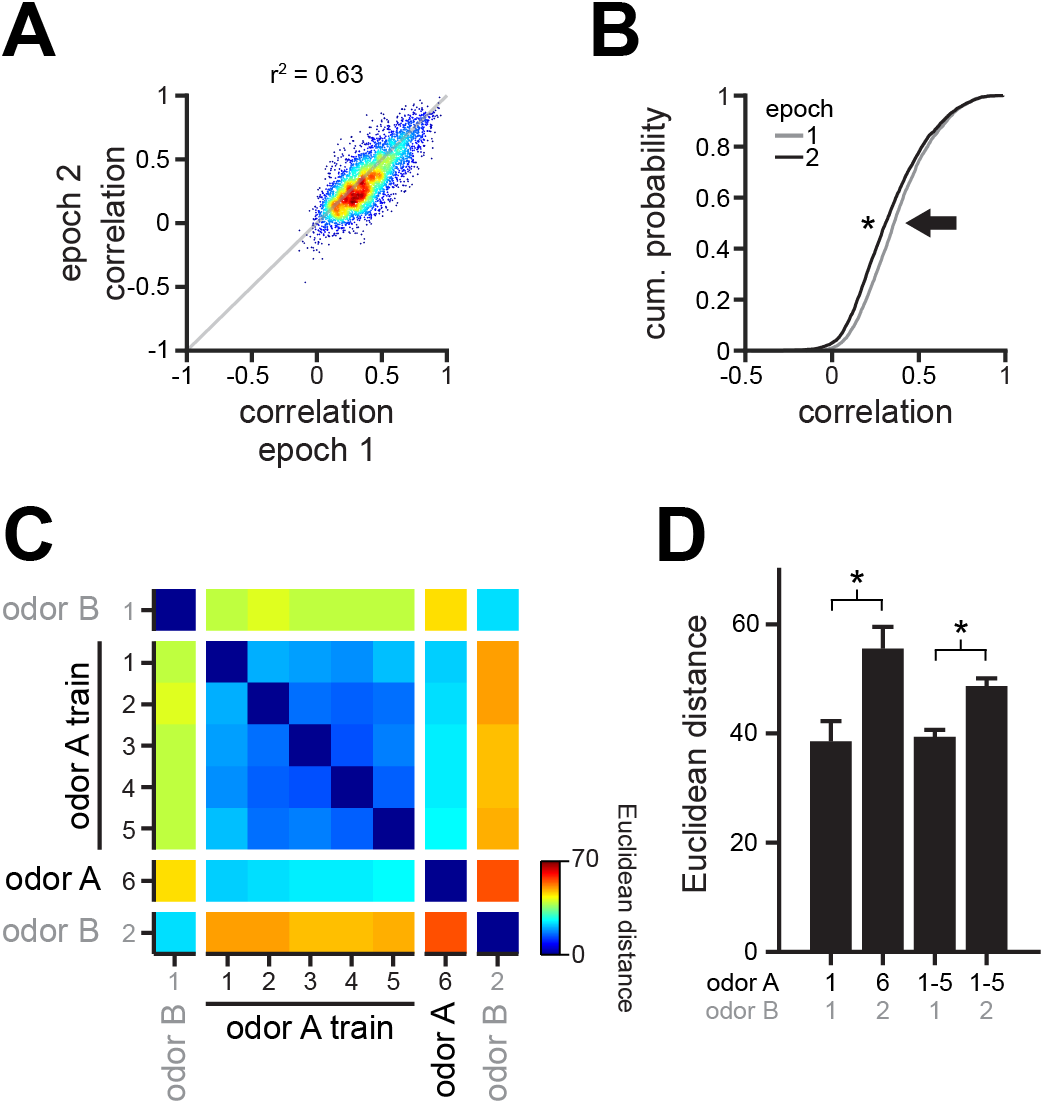
A. Correlation coefficients among all glomerulus pairs compared across the two ongoing activity epochs (n = 4,688 glomerulus pairs). B. Cumulative distribution of correlation coefficients for all recorded glomeruli. Correlation among glomeruli significantly decreases with repetitive stimulation (n = 309 glomeruli in 10 flies, Wilcoxon rank-sum test, p = 5.4 x 10-24). C. Matrix representing the average pairwise Euclidean distances among odor representations elicited by the unrepeated odor B and the repeated odor A, respectively (n = 10 flies). D. Euclidean distances among odor representations for the repeated odor A and unrepeated odor B. Note that the representation of the repeated odor A become more distant from that of the unrepeated odor B (mean ± s.e.m., n = 10 flies; Wilcoxon signed-sum test; odor A1-odor B1 vs odor A6-odor B2, p = 2.0 x 10-3; odor A1-5-odor B1 vs odor A1-5-odor B2, p = 0.02).

## Notes

### Competing Interest Statement

The authors have declared no competing interest.

### Summary of Updates

We corrected a single letter typo in the title

